# Duplication and overexpression of the genes encoding a beta 1,3-glucan synthase confer the intrinsic resistance to echinocandin in *Mucor circinelloides*

**DOI:** 10.1101/2022.01.03.474814

**Authors:** Alexis Garcia, Eun Young Huh, Soo Chan Lee

**Affiliations:** South Texas Center for Emerging Infectious Diseases (STCEID), Department of Molecular Microbiology and Immunology, The University of Texas at San Antonio, Texas, USA; Naval Research Unit – San Antonio (NAMRU-SA), Maxillofacial Injury and Disease Department

## Abstract

Procedures such as solid organ transplants and cancer treatments can leave many patients in an immunocompromised state resulting in an increased susceptibility to opportunistic diseases including fungal infections. Mucormycosis infections are continually emerging and pose a serious threat to immunocompromised patients. Currently there has been a sharp increase in mucormycosis cases as a secondary infection in patients battling SARS-CoV-2 infections. Mucorales fungi are notorious for presenting resistance to most antifungal drugs. The absence of effective means to treat these infections results in mortality rates approaching 100% in cases of disseminated infection. One of the most effective antifungal drug classes currently available are echinocandins. Echinocandins seem to be efficacious in treatment of many other fungal infections. Unfortunately, susceptibility testing has found that echinocandins have no to little effect on Mucorales. In this study, we found that the model Mucorales *Mucor circinelloides* genome carries three copies of the genes encoding for the echinocandin β-(1,3)-D-glucan synthase (*fks*A, *fksB*, and *fksC*). Interestingly, we revealed that exposing *M. circinelloides* to micafungin significantly increased the expression of the *fksA* and *fksB* genes when compared to an untreated control. We further uncovered that the serine/threonine phosphatase calcineurin is responsible for the overexpression of *fksA* and *fksB* as deletion of calcineurin results in a decrease in expression of all three *fks* genes and a lower minimal inhibitory concentration (MIC) to micafungin. Taken together, this study demonstrates that the *fks* gene duplication and overexpression by calcineurin contribute to the intrinsic resistance to echinocandins in *Mucor*.

**IMPORTANCE:** The recent emergence of mucormycosis cases in immunocompromised patients has become a more prevalent issue. The Mucorales fungi that cause these infections are known to be highly drug resistant, thus treatment options are limited and the mortalities of these types of infections remain unacceptably high. Echinocandins are one of the latest antifungal drugs developed to successfully treat fungal infections, but it remains unclear why Mucorales fungi are resistant to these treatments. In our study, we uncovered three copies of the genes (*fks*) encoding the drug target for echinocandins. Furthermore, we discovered that the serine-threonine phosphatase calcineurin is regulating the over-expression of these genes, which confers resistance. By inhibiting calcineurin we found that the expression of these drug targets decreases resulting in an increase in susceptibility to echinocandins both *in vitro* and *in vivo*

## INTRODUCTION

During the past decades, there has been an exponential growth in the discovery and implementation of advanced treatment for human diseases [1, 2]. This has led to better treatment for patients, and as a result we have been able to prolong human life. While these recent medical advances have certainly been beneficial overall, some have come with undesirable side effects. Procedures such as solid organ transplants and cancer treatments leave many patients in an immunocompromised state. This results in an increased susceptibility to emerging opportunistic diseases that flourish in immunosuppressed patients; including fungal infections that plague patients globally. Mucormycosis infections are continually emerging and pose a serious threat to immunocompromised patients [3–5]. Currently there has been a sharp increase in mucormycosis cases as a secondary infection in patients battling SARS-CoV-2 infections [6, 7]. Fungi in the order Mucorales are etiologic agents of mucormycosis and notorious for presenting resistance to most antifungal drugs [8, 9]. Once mucormycosis has been diagnosed in a patient, treatment guidelines suggest that physicians begin with surgical debridement of the infected and necrotic tissue followed by treatments with a first line antifungal drug such as lipid complex amphotericin B [10]. Amphotericin B is known to be a highly nephrotoxic compound posing an additional threat to many patients, particularly those presenting with renal related complications [10, 11]. In the event a patient is not able to withstand treatment by amphotericin B, salvage therapy involving treatment with an azole such as posaconazole or Isavuconazole is recommended, despite the azoles having a limited treatment efficacy [10, 12]. More importantly, Mucorales fungi exhibit intrinsic short-chain azole resistance [13, 14]. These less-than-ideal treatment options are the only options currently available for patients. Thus, in the absence of effective means to treat such infections, mortality rates associated with a disseminated mucormycosis case approach 100% [15, 16].

One of the most effective antifungal drug classes currently available for patients are echinocandins [17]. Echinocandins are a group of semisynthetic lipopeptides harboring an N-linked acyl lipid side chain that act as non-competitive inhibitors of the β-(1,3)-D-glucan synthase, a key component necessary for the integrity of the β-(1,3)-D-glucan synthase enzyme is encoded by the *fks* gene family, which is conserved across fungi [18]. The known function of β-(1,3)-D-glucan, which is then incorporated into the fungal cell wall to maintain its structural integrity [18]. Echinocandins are known to impart an excellent fungicidal activity against *Candida albicans* and some activity against *Candida glabrata* [20, 21]. Although echinocandins seem to be efficacious in treatment of other fungal infections, previous susceptibility testing has found that echinocandins have little to no effect on Mucorales fungi [22–24]. However, the mechanisms responsible for this resistance of Mucorales fungi to echinocandins remain unclear [25]. Like other fungi, Mucorales possess a cell wall which is the normal site of action of echinocandins, yet these drugs show no effect when used to treat mucormycosis infections. To elucidate the intrinsic resistance mechanisms in Mucorales, we investigate the model Mucorales *Mucor circinelloides* (denoted *Mucor*). In our study, we found that the *Mucor* genome carries three copies of the *fks* gene in its genome (*fksA*, *fksB*, and *fksC*). A phylogenetic analysis of the *fks* genes revealed that *fksA* and *fksB* were converged from a segmental duplication in an early divergence of Mucorales and *fksC* is distinctly related to the other two. Unlike what we observe in other fungal organisms, the *fksA* and *fksB* genes were found to be essential. In our study we were able to obtain heterokaryotic mutants of the *fksA* and *fksB* genes and a homokaryotic mutant of the *fksC* gene. We found that when *Mucor* is exposed to micafungin, the expressions of the *fksA* and *fksB* genes are elevated 15-fold when compared to an untreated control while *fksC* was down regulated. We further revealed that the serine/threonine phosphatase calcineurin is responsible for the overexpression of *fksA* and *fksB* as deletion of calcineurin results in a decrease in the expression of all three *fks* genes and a lower minimal inhibitory concentration (MIC) to micafungin. In addition, deletion of the calcineurin regulatory gene *cnbR* results in a lower MIC to micafungin and a higher *in vivo* susceptibility to micafungin compared to wild-type (WT); on the other hand, the *fksC*Δ mutants exhibit a higher MIC and lower susceptibility similar to the WT. Taken together, this study demonstrates that the *fks* gene overexpression by calcineurin contribute to the intrinsic resistance to echinocandin in *Mucor*.

## RESULTS

### Identification of fks genes in Mucor: duplication of the fks genes

The *Saccharomyces cerevisiae fks1* gene was used to identify *fks* genes in the *Mucor* genome. BLAST searches revealed that there are three copies of *fks* genes in *Mucor*: *fksA* (encoding OAD02999), *fksB* (OAD08543), and *fksC* (OAD00805). We examined the molecular phylogeny of the three *fks* genes in *Mucor* and two other Mucorales *Rhizopus delemar* and *Phycomyces blakesleeanus* and found that *fksA* and *fksB* share a common ancestral gene that diverged prior to species differentiation and generated precursors *fksA* and *fksB*, which eventually resulted in the differentiation of *fksA* and *fksB* (Fig. 1). Furthermore, *fksC* seems to have diverged prior to this event resulting in *fksC* to not be as closely related to *fksA* and *fksB*, something that is observed in other fungi with two distinctly related *fks* genes. The same phylogenetic patterns can be observed in other Mucorales fungi such as *Phycomycetes blakesleeanus* and *Rhizopus delemar* (Fig. 1). In *C. glabrata* and *S. cerevisiae* there are also three *fks genes* (*fks1, fks2,* and *fks3*), similar to what we observe in *Mucor* (Fig. 1); however, the *fks* gene duplications in Mucorales appear to be more ancestral and those are from a more recent whole genome duplication in the *Saccharomyces sensu stricto* [26]. The FksA and FksB share a 43.97% identity, while FksA vs FksC or FksB vs FksC at 40.17% and 31.65%, respectively.

**Figure 1.**
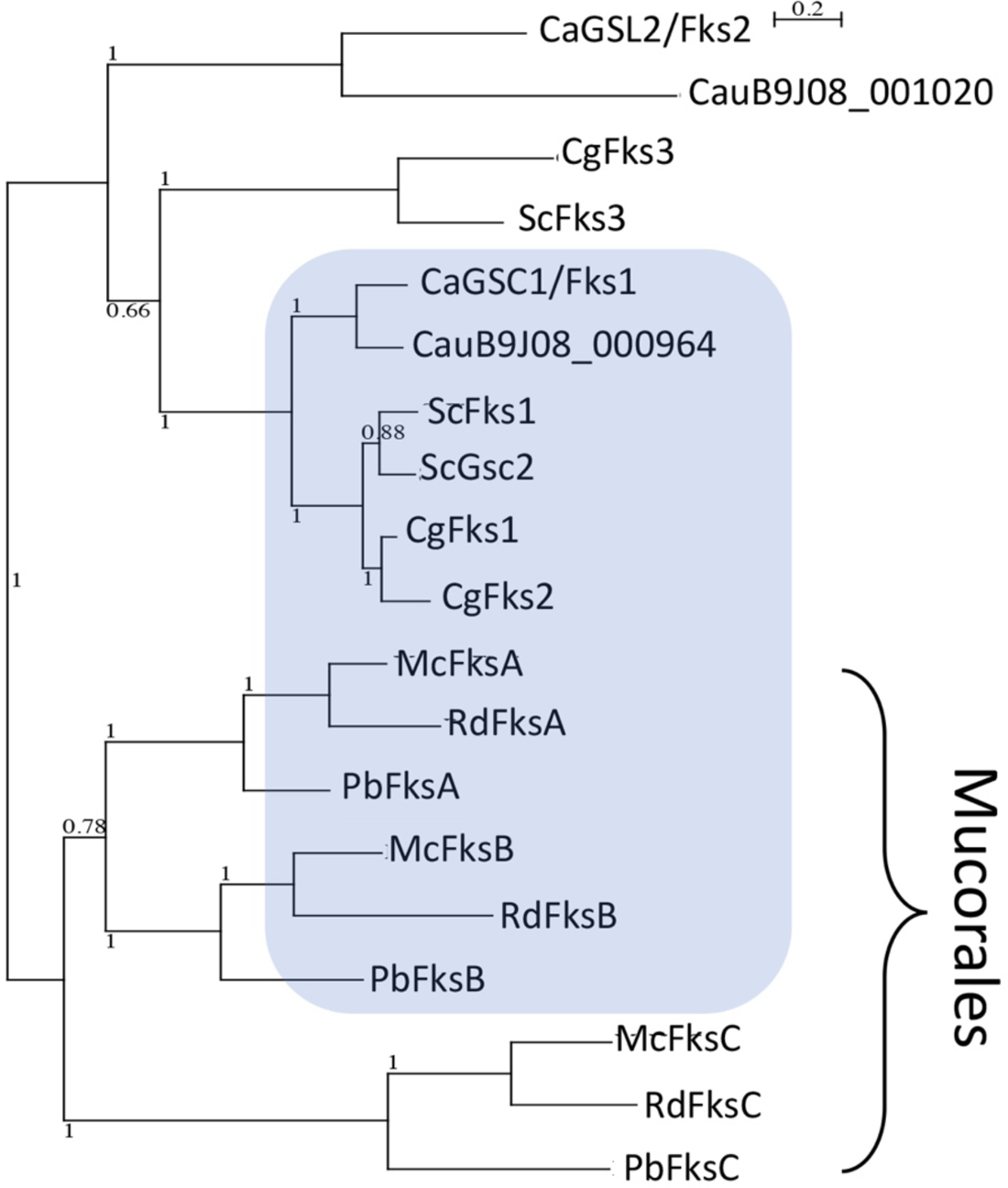
Phylogeny of Fks proteins. The shading indicates major Fks proteins. Ca = *C. albicans*, Cau = *C. auris*, Cg = *C. glabrata*, Sc = *S. cerevisiae*, Mc = *M. circinelloides*, Pb = *P. bkalesleeanus*, Rd = *R. delemar*. Scale indicates an amino acid substitution per position. An ancestral fks gene may have diverged before divergence between Mucoromycota and Ascomycota. Before speciation in Mucorales, the common ancestral *fks* gene appears to have been diverged into two different paralogs, *fksC* and a common ancestor of *fksA* and *fksB*, which later diverged into *fksA* and *fksB*.

The *Mucor* genomes conserve the target *fks* genes encoding a β-1,3-glucan synthase; however, it exhibits a higher level of resistance. To understand the mechanisms of the resistance, we further investigate the functions of the *fks* genes.

### Overexpression of *fks* genes in response to micafungin in *Mucor*

In *C. glabrata,* the β-(1,3)-D-glucan synthase products of all three *fks* genes are functionally redundant. The *fks1* and *fks2* genes show a similar expression pattern marked by up-regulation during most of the early life of the fungus, whereas the *fks3* gene is down-regulated until the fungal sporulation stage [26]. We therefore hypothesize that a similar relationship exists among the *fks* genes (*fksA, fksB* and *fksC*) found in *Mucor*, indicative of a similar functional redundancy between the forms of β glucan synthase encoded by *fksA* and *fksB*, plus unique expression of *fksC* during the different developmental stage. However, when we measured the expression of the *fks* genes in *Mucor* using quantitative real-time PCR (qRT-PCR), we did not observe a differential expression pattern of the three *fks* genes in the different developmental stages, including isotropic growth phase of spores, germination phase, and sporulation phages (Fig 1S).

We further measured the expression of the *fks* genes to determine the expression profile of the *Mucor fks* genes during the two clinically used echinocandins micafungin and caspofungin challenge (Fig. 2). Interestingly, we observed a significant up-regulation 15-fold (p=0.0055) of the *fksA* and 13-fold (p=0.0026) *fksB* transcripts when *Mucor* grew in the presence of micafungin (50 μg/ml) compared to an unchallenged control (Fig. 2A). However, the expression of *fksC* was downregulated (p=0.0045). This demonstrates that the elevated expression of the β synthase-expressing genes is affected by echinocandins, which can result in resistance to the drug. It is possible that *fksA* and *fksB* share a major functional similarity in the β-(1,3)-D-glucan synthase. Interestingly, caspofungin (10 μg/ml) treatment did not elevate the expression of *fks* genes (Fig 2B). Indeed, the presence of caspofungin resulted in lowered expression of the *fksB* and *fksC* genes, whereas that of *fksA* did not change.

**Figure 2.**
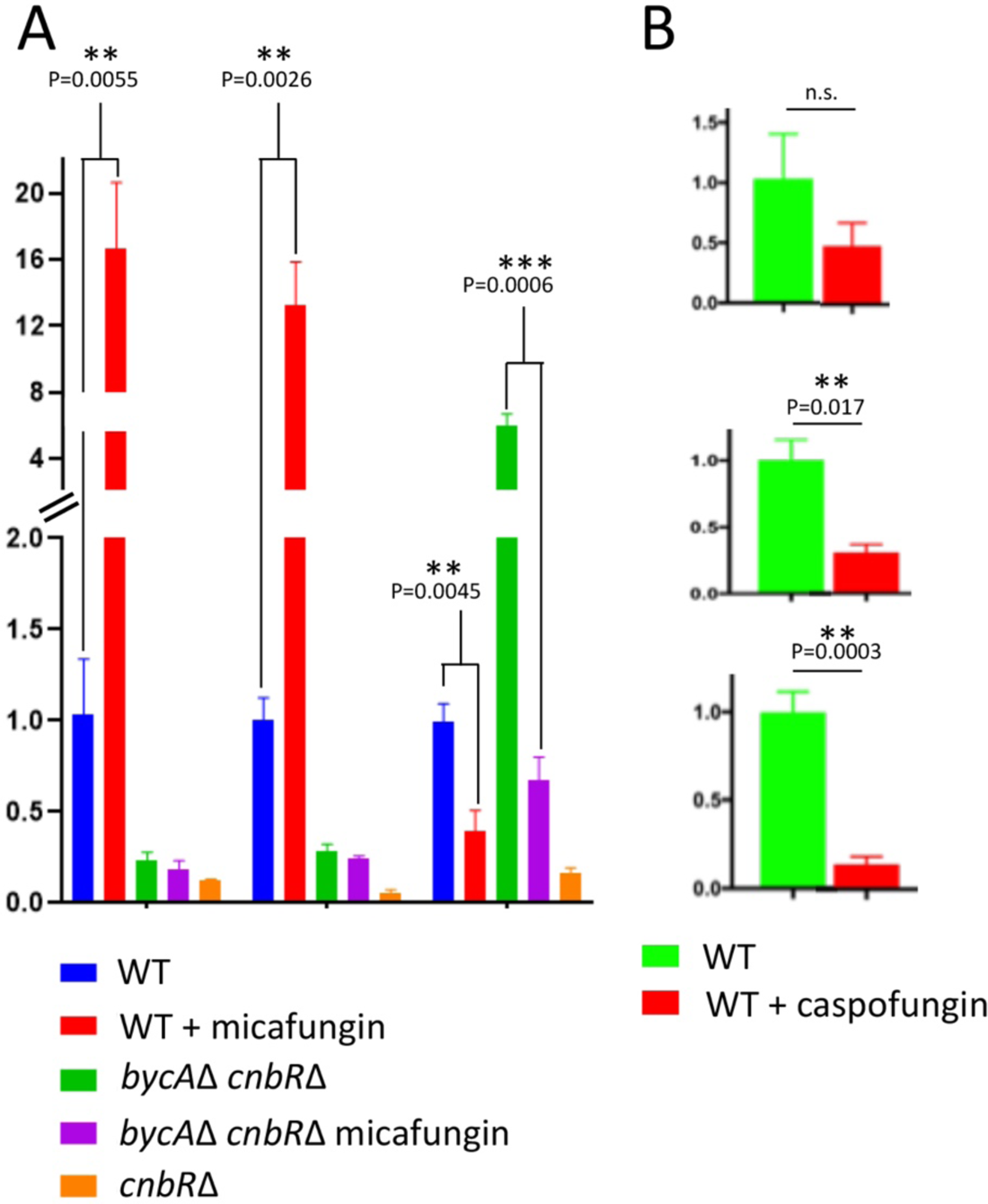
Expression of the three fks genes in Mucor. (A) the expression of *fks* genes in the presence of micafungin. In the presence of micafungin the *fksA* and *fksB* genes are overexpressed 16-fold and 13-fold respectively. Mutants lacking a functional calcineurin present an overall down regulation of the *fks* genes even in the presence of micafungin. (B) The expression of the *fks* genes in the presence of caspofungin. In the presence of caspofungin *fksA* gene had no difference in expression while the *fksB* and *fksC* genes were down regulated significantly.

### Intrinsic resistance to micafungin by overexpression of *fks* genes regulated by calcineurin

In *S. cerevisiae*, the expression of *FKS2* but not *FKS1* is induced by calcineurin [27] and in *C. glabrata* calcineurin participate in a transcriptional regulation of *FKS2* but not *FKS1* [28]. In *Mucor*, our previous study verified that the calcineurin inhibitor FK506 exhibits an in vitro synergistic effect with micafungin [29]. These previous observations prompted us to test whether calcineurin is involved in the overexpression of the *fks* genes in *Mucor*. We therefore measured the expression of *Mucor* mutants lacking a functional calcineurin.

The *Mucor* genome encodes a calcineurin regulatory B subunit (CnbR), and the deletion of the *cnbR* gene results in a complete loss of calcineurin function. The *cnbR*Δ mutants are locked in the yeast phase [30, 31], which is not a normal growth status of the fungus as the cell wall components of hypha and yeast are different (data not shown). To test the susceptibility to micafungin and its relationship to calcineurin, we added *cnbR*Δ *bycA*Δ double mutants with *bycA*Δ as a control, in which the secondary mutation in the *bycA* gene encoding an amino acid permease enables the *cnbR*Δ mutants to grow in a hyphal form. Therefore, we can evaluate the expression of *fks* genes in a mold form in the presence or absence of roles of calcineurin function (Fig. 2A). Interestingly, in both the calcineurin mutants *cnbR*Δ and *cnbR*Δ *bycA*Δ, the expression of *fksA* and *fksB* genes were significantly lowered regardless of the presence of micafungin compared to the wild type and *bycA*Δ, respectively (Fig. 2A).

To measure the extent to which each of the different mutants we generated impacts the susceptibility of *Mucor* to micafungin or caspofungin, we employed a minimum inhibitory concentration (MIC) assay. In brief, we inoculated spores of wild-Δ::*cnbR* in RPMI media containing micafungin or caspofungin serially diluted in twelve equal steps ranging from of 256 µg/ml to 0 µg/ml. Minimum inhibitory concentrations were determined according to the Clinical and Laboratory Standards Institute (CLSI) guideline (Fig 3) [32].

**Figure 3.**
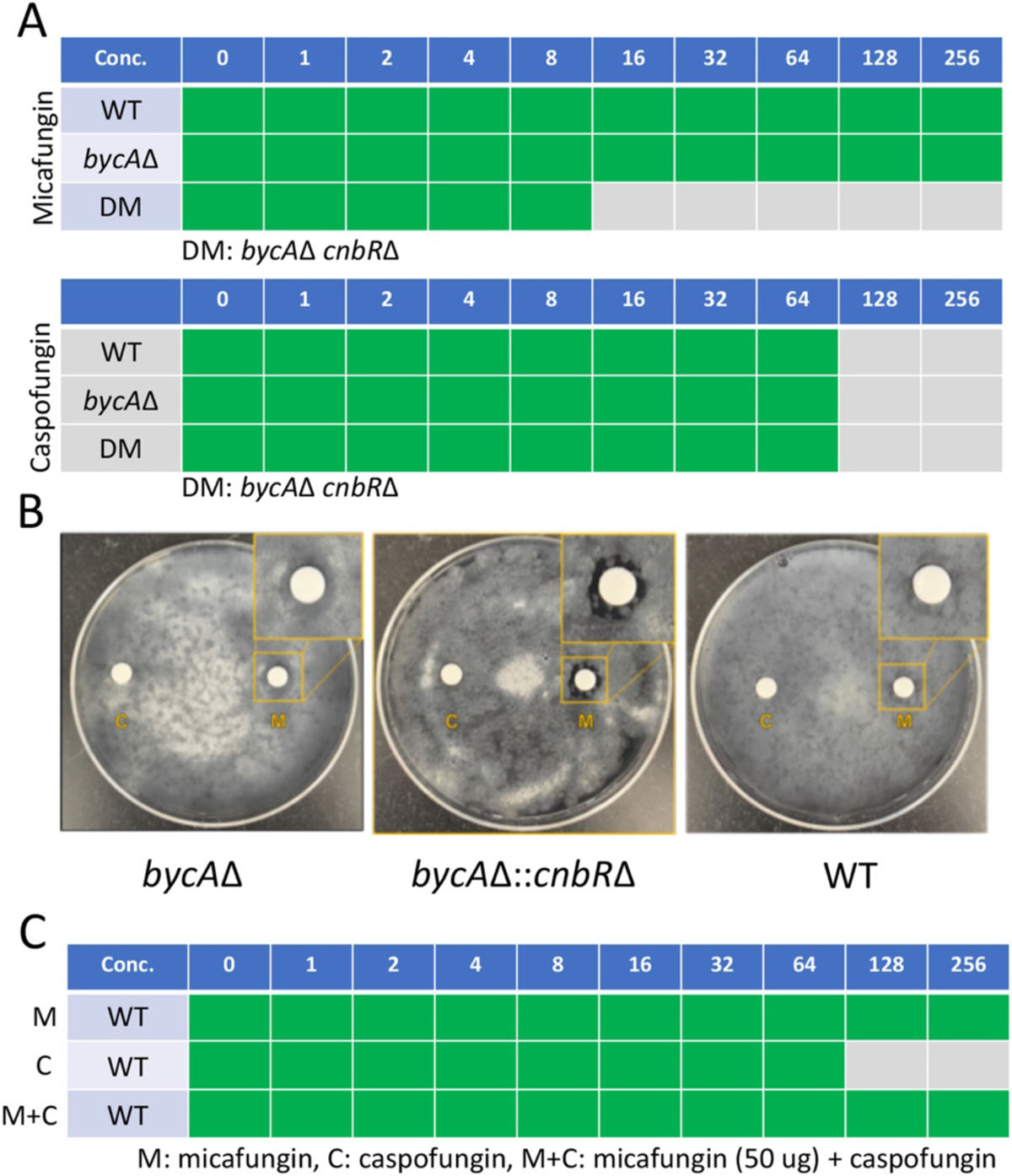
Drug sensitivity between caspofungin and micafungin is different in the wild-type (WT) strain. (A) Minimum inhibitory concentration assay (MIC) reveals sensitivity to micafungin in the calcineurin mutant *cnbR*Δ *bycA*Δ (DM). Deletion of the calcineurin regulatory subunit gene *cnbR* and the amino acid permease gene *bycA* results in a lowered MIC of 16 μg/ml compared to that of wild-type or *bycA*Δ. Additionally, the MIC of caspofungin was unanimously 128 μg/ml across the tested strains. (B) Disk diffusion assay confirms sensitivity to micafungin (M) and not caspofungin (C). Paper disks loaded with 100μg of either antifungal drug was placed on lawns of either the WT strain, *bycA*Δ mutant or DM. a zone of inhibition around the disk loaded with micafungin was observed in the DM strain, but not in any of the other strains tested. (C) A combination of both micafungin and caspofungin causes an increase in the MIC of the WT strain. When micafungin is solely present the MIC of the WT strain is >256 μg/ml. when caspofungin is solely present the MIC of the WT strain is 128 μg/ml. When both drugs are administered in combination the MIC of the WT strain is increased to >256 μg/ml. This recapitulates our findings that the overexpression of the *fksA* and *fksB* genes confers resistance to echinocandins.

The calcineurin mutant *cnbR*Δ *bycA*Δ was more sensitive to micafungin with a MIC of 16 g/ml compared to that of wild-type or *bycA* (Fig 3A). The MIC of these mutants to caspofungin were observed to be lower at 128 μg/ml for both wild type and calcineurin mutants (Fig 3A). In addition, a disc diffusion assay clearly demonstrates that the calcineurin mutant *cnbR*Δ *bycA*Δ was more sensitive compared to the two controls, wild-type and the *bycA* mutant (Fig 3B). The *Mucor* genome encodes three copies of calcineurin catalytic A subunits, CnaA, CnaB, and CnaC. To test if each catalytic subunit is differentially associated with the echinocandin resistance, we further deleted *cnaC* (Fig 2S) in addition to *cnaA* and *cnaB* that we have disrupted in previous studies [30]. However, each *cna* mutants did not exhibit difference in susceptibility to micafungin or caspofungin (Fig 3S).

*Mucor* responds to micafungin and caspofungin differently, in which micafungin elevates the expression of *fskA* and *fksB* but caspofungin does not (Fig 1).These difference responses are correlated with different MIC’s to the two drugs, in which MIC to micafungin is >256 (μg/ml) and that to caspofungin is 128 (Fig 3C). When *Mucor* was challenged with caspofungin in the presence of micafungin (50 μg/ml), the MIC to caspofugin was elevated to >256 (Fig 3C). Overexpression of fksA and fksB caused by micafungin results in resistance to caspofungin. This result further confirms that the overexpression of the *fksA* and *fksB* genes confer resistance to echinocandins.

These results demonstrate that calcineurin regulates the overexpression of the *fksA* and *fksB* genes to confer natural resistance to micafungin in *Mucor*. Echinocandin resistance mechanism by the overexpression of the target genes is novel and has not previously been reported in any fungal systems.

### Deletion of the *fksA* or *fksB* genes results in an elevated sensitivity to echinocandins

In *Candida* species, the *FKS1* play major roles with a redundant role of the *FKS2* gene when the *FKS1* gene is deleted. In *Mucor*, it is possible that *fksA* and *fksB* play a more important role than *fksC* based on the observation that *fksA* and *fksB* are upregulated in the presence of micafungin, but *fksC* is not. In addition, when the *fksC* gene was deleted and the deletion was confirmed by junction PCRs (Fig S4) and Southern blot (data not shown), the MIC of and sensitivity to micafungin of *fksC*Δ mutants are not different from that of wild-type (> 256 μg/ml) (Table 1). This indicates the *fksC* is a minor *fks* gene in *Mucor*.

**Table 1.** MICs of *Mucor fks* mutant strains to caspofungin and echinocandin.

| Strain | Caspofungin ($\mu\text{g/ml}$) | Micafungin ($\mu\text{g/ml}$) |
| --- | --- | --- |
| WT | 128 | >256 |
| *fksA*$\Delta^{wt}$ | 128 | >256 |
| *fksB*$\Delta^{wt}$ | 128 | >256 |
| *fksC*Δ | 128 | >256 |

We then attempted to delete *fksA* and *fksB* by utilizing our previously developed recyclable marker system [33, 34] and obtained hetorogolous mutants at each gene, where the deletions were comfirmed by junction PCRs and (Fig 5S and 6S, respectively) and Southern blot (data not shown). Although we produced mutant alleles lacking either *fksA* or *fksB,* we were unable to obtain homokaryotic *fksA*Δ or *fksB*Δ mutants through intensive vegetative selections; instead, the mutants always existed as heterokaryons that carried both the *fksA* and *fksA*Δ::*pyrG-247* or *fksB* and *fksB*Δ::*pyrG-247* alleles. Therefore, the mutants denote as *fksA*^wt/Δ^ or *fksB*^wt/Δ^, respectively.

Gene expression was measured via quantitative real-time PCR for the *fksA* and *fksB* genes in *Mucor* during echinocandin challenge (Mycamine 50µg/ml) in the WT, *fksA*^wt/Δ^, *fksB*^wt/Δ^, and in an untreated control (Fig 4). The partial deletion mutants resulted in a lowered expression of their respective *fks* gene. We observed a significant down regulation of the *fksA* gene in the *fksA*^wt/Δ^ mutant when compared to the WT control. A similar result could be observed in the expression of the *fksB* gene that was also significantly down regulated in the *fksB*^wt/Δ^ mutant. In a test of MICs of each mutant to echinocandins, we did not observe differences between groups (Table 1). It is possible that growth inhibition of the partial deletion mutants by the drugs on a 96 well plate might not be as effective considering the mutants still express the WT *fks* alleles responsible for the resistance to echinocandins. To circumvent this, we utilized a disk diffusion assay to observe a zone of inhibition around paper disks (Fig 4B). In brief, we plate a lawn of either WT, *fksC*, *fksA*^wt/Δ^, or *fksB*^wt/Δ^. We then place three disks loaded with micafungin (100 µg). In these experiments, we were able to observe the zones of growth inhibition of the *fksA*^wt/Δ^ and *fksB*^wt/Δ^ mutants. On the other hand, a zone of inhibition was not observed in the WT or the *fksC* mutant. These results can be an analogy of haplo-insufficiency and importantly demonstrate that a gene duplication of the fks genes into *fksA* and *fksB* in *Mucor* confer resistance to echinocandins

**Figure 4.**
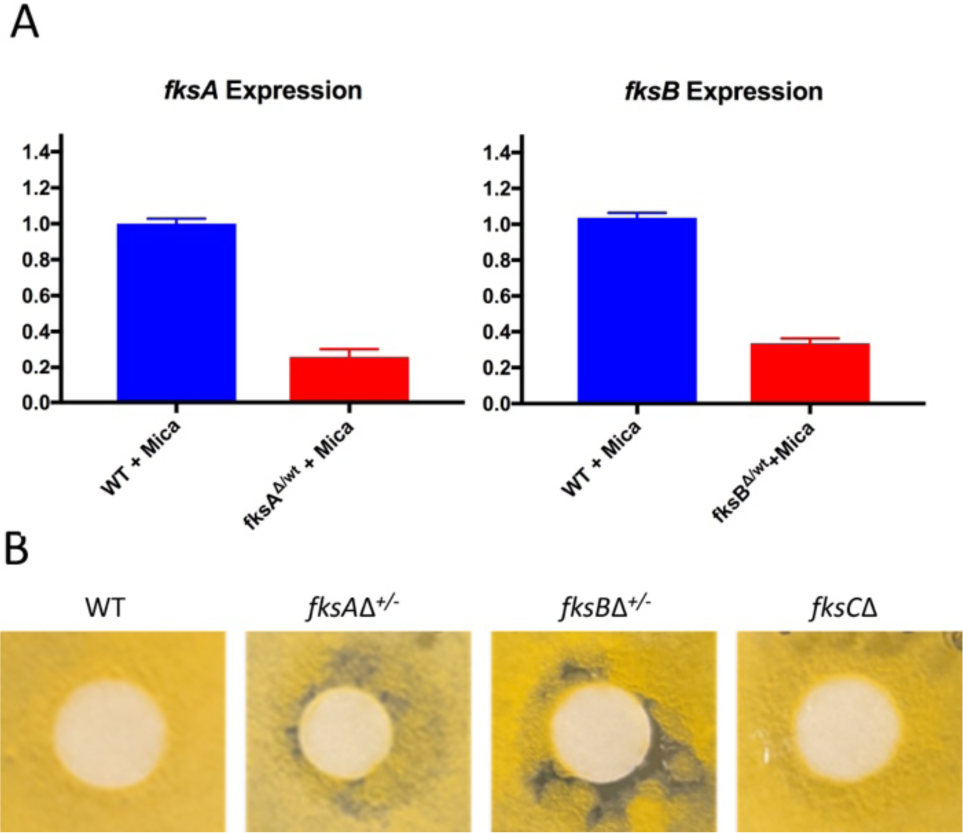
Deletion of *fksA* or *fksB* results in an increased susceptibility to micafungin. (A) Expression of the *fksA* gene in the *fksA*^wt/Δ^ mutant and the *fksB* in the *fksB*^wt/Δ^ were measured and compared to the expression of each respective gene in a wild-type (WT) control treated with 50 µg/ml of micafungin. Expression of the *fksA* gene was lower in the *fksA*^wt/Δ^ mutant in the presence of micafungin when compared to a WT control. The expression of the *fksB* gene was also lower in the *fksB*^wt/Δ^ mutant in the presence of micafungin. (B) Disk diffusion assay reveals increased sensitivity to micafungin in the *fksA*^wt/Δ^ and *fksB*^wt/Δ^ mutants but not the *fksC* mutant. Paper disks containing 100 µg of micafungin were placed on a lawn of spores spread on MMC (pH 4.5) media. Zones of inhibition can be observed in the *fksA*^wt/Δ^ and *fksB*^wt/Δ^ mutants but not the *fksC* mutant or WT control.

### Mutants lacking a functional calcineurin are more susceptible to micafungin in a heterologous mucormycosis model

Previously we observed the MIC of the *bycA*Δ *cnbR*Δ to micafungin was lower than our WT. As previously mentioned, mutants lacking a functional calcineurin had a lowered expression of the *fksA* and *fksB* genes. To further test our hypothesis we moved on to an *in vivo* model. We injected galleria mellonella larvae (*n*=15/group) with 40,000 spores of the *bycA*Δ *cnbR*Δ mutant or with phosphate-buffered saline (PBS). Three groups were treated with 16 mg/kg of micafungin and three groups remained untreated. We then monitored survival for 10 days. The treated groups exhibited an increased survival rate when compared to the untreated controls (Fig 5).

**Figure 5.**
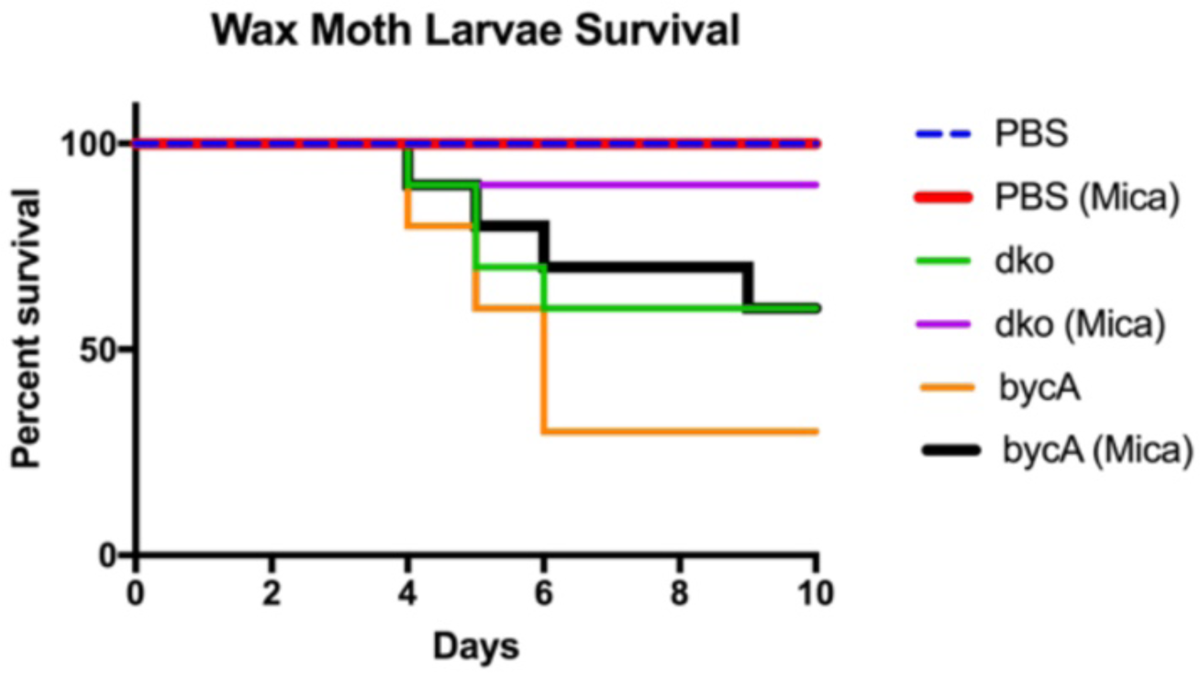
Calcineurin mutants were more susceptible to micafungin treatment in a *Galleria mellonella* (wax moth) model of mucormycosis. Wax moth larvae were inoculated with 4×10^4^ *Mucor* spores in a 2 µl mixture containing PBS. The spores were administered via injection on the last left proleg. The larvae were monitored for survival over the course of ten days.

## DISCUSSION

Echinocandins are some of the most effective antifungal drug classes currently available for patients [17]. In clinical settings, they lead to the efficacious outcome in the treatment of other fungal infections including candidiasis infections and invasive aspergillosis [35, 36]. Previous studies have found that a combination therapy of caspofungin with amphotericin B lipid complex can result in a synergistic effect for mucormycosis [37]. In clinical sets, echinocandins are used to treat patients with mucormycosis although the efficacy is in a question due to Mucorales fungi being highly resistant to echinocandins. Despite the Mucorales genomes encoding for the target β-(1,3)-D-glucan synthase enzyme like many other fungi, it has yet to be elucidated how Mucorales fungi achieve the high level of intrinsic resistance to these drugs.

Acquired resistance mechanisms to echinocandin in *Candida* spp have been reported [38]. Most cases involve genetic mutations in the frequently mutated regions identified as hotspots of the *fks* genes. In *C. albicans* there are three known hot spot regions that can result in echinocandin resistance if there are mutations within these regions.

Our study demonstrates that one of the mechanisms for intrinsic resistance to echinocandin in Mucorales is overexpression of the target of the drug, in which *Mucor* overexpresses the *fksA* and *fksB* genes in the presence of echinocandins. There are reports that the overexpression of the *erg* genes can result in azole resistance [39]. Thus, it is possible that the *fks* gene overexpression observed in *Mucor* is involved in echinocandin resistance. In addition, the presence of two copies of the major *fks* genes (*fksA* and *fksB*) can also contribute to the resistance of echinocandin treatment.

In *C. albicans* and *S. cerevisiae*, calcineurin regulates the expression of *FKS2* gene via the transcription factor Crz1. Interestingly, the paralog *FKS1* gene is not regulated by calcineurin. In *Mucor*, however, calcineurin regulates both of the *fksA* and *fksB* genes and this regulation results in highly elevated expression of the *fks* genes unlike in two yeast species. Similarly in *Cryptococcus neoformans* the underlying mechanisms for echinocandin resistance has been isolated to the involvement of Cdc50, a regulatory subunit of a lipid flippase [40]. Previous studies have found that Crm1, a known homolog of mechanosensitive channel proteins, can promote caspofungin resistance via a heightened activation of the calcineurin pathway [40]. In addition, in *C. glabrata*, the two *FKS1* and *FKS2* genes are known to be redundant and expressed in different developmental stages. However, in *Mucor*, as we were not able to generate the complete deletion of either of *fksA* or *fksB* gene, it is possible that the genes are not completely redundant and play their own essential roles. These differences imply an independent and unique evolutionary trajectory in the regulation of *fks* genes by calcineurin in the two different groups of fungi.

Calcineurin is required for invasive hyphal growth in *Aspergillus fumigatus*, for growth at 37°C in *Cryptococcus neoformans*, and for serum tolerance in *Candida albicans* [41–43]. Calcineurin, therefore, has long been considered a promising drug target. When *Mucor* is grown in the presence of the calcineurin inhibitor FK506, it exhibits yeast growth instead of hyphae, which shows that calcineurin is required for invasive hyphal growth in this fungus (yeast to hyphal transition) [31]. Mutants lacking functional calcineurin are less virulent than the wild type, which suggests that calcineurin is a key target in the treatment of mucormycosis [30, 31, 44]. In addition to the calcineurin inhibitors’ activity, our study provides a novel insight how to improve an already existing drug. Targeting calcineurin in *Mucor* can indeed sensitize the fungal cells to become susceptible to micafungin.

How does calcineurin up-regulate the *fks* genes in *Mucor*? Ortholog of the transcription factor Crz1 found in *S. cerevisiae*, *Candida* spp. and *C. neoformans* [45–47] are identified in the *Mucor* genome. As Crz1 is diverged in sequences, a BLAST analysis may not result in a Crz1 homolog in Mucorales. As Crz1 is dephosphorylated by calcineurin, a comparative phosphoproteomics and functional study will be required to identify a Crz1 ortholog in *Mucor* that plays a key role in the echinocandin resistance. This study is on-going in our group to identify the key transcription factor.

Calcineurin plays essential roles in virulence in many pathogenic fungi and therefore has been targeted to develop novel antifungal drugs [48, 49]. Our study further support that calcineurin inhibitors can be used in combination with echinocandins as they enforce Mucorales more sensitive to echinocandins. However, the inhibition of calcineurin function by gene knock outs or drugs can make *Mucor* more sensitive to micafungin. However, the MIC to echinocandins of the calcineurin mutants is still significantly higher compared to that in *C. albicans*, for example, it’s 0.008 µg/ml in *C. albicans* and 16 µg/ml in *Mucor*. Even acquired echinocandin resistant *C. albicans* isolates have an MIC of >1 µg/ml. Due to this high level of MICs in *Mucor*, we did not observe efficacy in a murine mucormycosis model with a monotherapy with micafungin (data not shown) although the wax moth mucormycosis model exhibited micafungin potentially rescues the infected larvae (Fig 5). Therefore it is possible that the FksA and/or FksB themselves are resistant to echinocandins, which our studies are currently on-going to address this question. Indeed, in a preliminary study we found that *Mucor* presents intrinsic differences in these hotspots when we align the sequences of *C. albicans* hotspot 1 with the hotspot 1 region in *Mucor*. It is possible that the β glucan synthase enzyme in *Mucor* is conferring an intrinsic resistance to echinocandins by naturally expressing a hotspot region that is known to be resistant to echinocandins.

## MATERIALS AND METHODS

### Ethics statement

The animal experiments in this study were conducted at the University of Texas at San Antonio in accordance with the institutional Animal Care and Use Committee (IACUC) guidelines and in full compliance with the United States Animal Welfare (Public Law 98-198) and National Institute of Health (NIH) guidelines. The animal protocol MU104 used in this study was approved by the UTSA IACUC. The experiments were conducted in a Division of Laboratory Animal Resources (DLAR) facility, which is accredited by the Association for Assessment and Accreditation of Laboratory Animal Care (AAALAC).

### Strains and culture conditions

Strains and plasmids used in this study is listed in Table 2.

**Table 2.** Strains used in this study

| Strain | Genotype | Remark |
| --- | --- | --- |
| CBS277.49 | Wild-type *Mucor circinelloides* |  |
| R7B | CBS277.49 origin, *leuA*<sup>-</sup> |  |
| MU402 | R7B origin, *leuA*<sup>-</sup> *pyrG*<sup>-</sup> |  |
| MSL7 | MU402 origin, *cnbRΔ::pyrG leuA*<sup>-</sup> | Lee et al 2013 |
| MSL45 | MU402 origin, *bycAΔ::pyrG-dpl237 leuA*<sup>-</sup> | Vellanki et al 2020 |
| MSL68 | MSL45 origin, *cnbRΔ::leuA bycAΔ::pyrG-dpl237* | Vellanki et al 2020 |
| MSL9 | MU402 origin, *cnaAΔ::pyrG leuA*<sup>-</sup> | Lee et al 2013 |
| MSL22 | MU402 origin, *cnaBΔ::pyrG leuA*<sup>-</sup> | Lee et al 2015 |
| MSL35 | MU402 origin, *cnaCΔ::pyrG-dpl237 leuA*<sup>-</sup> | This study |
| MSL36 | MU402 origin, *cnaCΔ::pyrG-dpl237 leuA*<sup>-</sup> | This study |
| MAG11 | MU402 origin, *fksCΔ::pyrG-dpl237 leuA*<sup>-</sup> | This study |
| MAG12 | MU402 origin, *fksCΔ::pyrG-dpl237 leuA*<sup>-</sup> | This study |
|  |  | This study |
| MAG13 | MU402 origin, *fksA*<sup>wt/Δ</sup>, harboring *fksA::pyrG-dpl237 leuA*<sup>-</sup> |  |
| MAG14 | Same as above |  |
|  |  | This study |
| MAG15 | MU402 origin, *fksB*<sup>wt/Δ</sup>, harboring *fksA::pyrG-dpl237 leuA*<sup>-</sup> |  |
| MAG16 | Same as above |  |

**Table 3.**
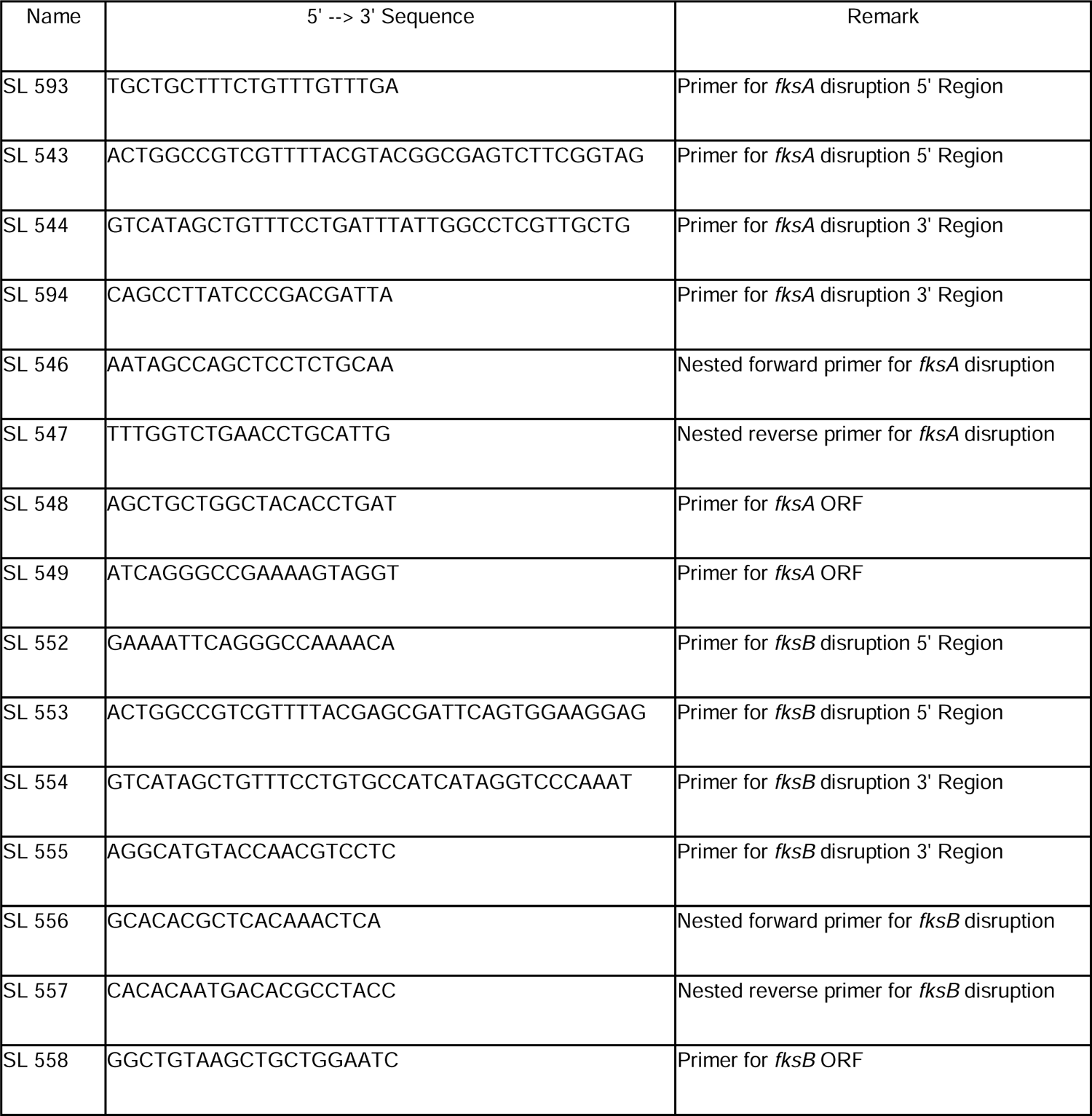
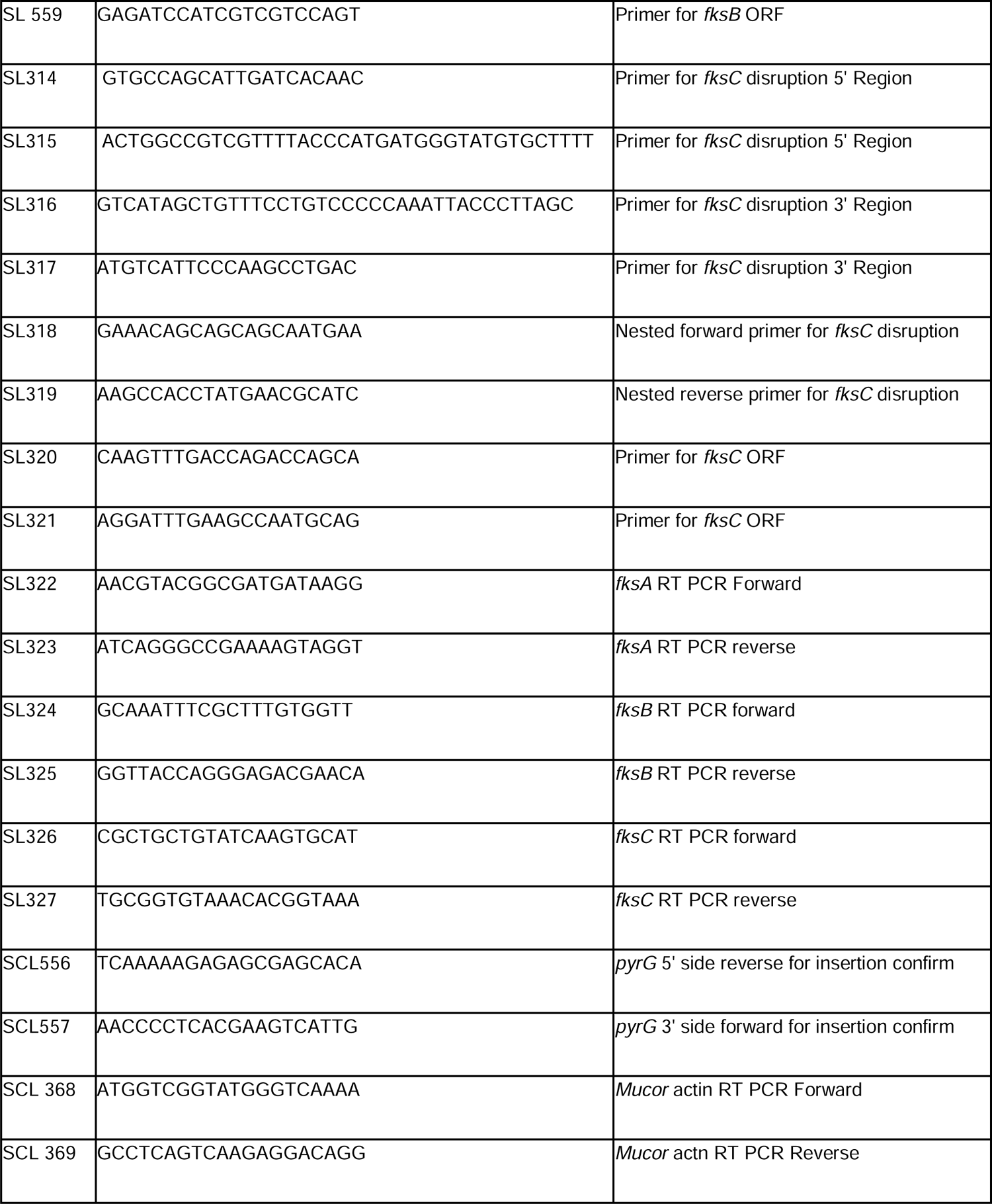
Primers used in this study.

*Mucor* strains were grown for propagation at 26°C in minimal media with casamino acids (MMC) at a pH of 4.5. Colony isolation was achieved by growing our Mucorales strains on MMC media at a pH of 3.2, which results in the formation of clustered colonies. To evaluate the minimum inhibitory concentration of echinocandins; Mucorales strains were grown in 96 well plates containing RPMI media which was treated with serially diluted echinocandins ranging from 256 µg/ml to 0.25 µg/ml. The 96 well plate were incubated at 26°C. Inhibition of growth was evaluated via microscopy.

### Quantitative PCR for *fks* gene

The RNAs from *Mucor* strains were extracted using the MasterPure™ yeast RNA purification kit (Lucigen MPY03100). Extracted RNA was then converted to cDNA by utilizing a high-capacity cDNA reverse transcription kit (Applied Biosystems™ 4374966). Quantitative real-time PCR was then conducted on (MACHINE INFO GOES HERE) using the PowerUp™ SYBR™ green master mix (Applied Biosystems™ A25742). To amplify the *fksA* gene we utilized the SL322 and SL323 primers; SL324 and 325 were used to amplify *fksB*. Finally, *fksC* was amplified with the primers SL326 and SL327. As a control we amplified the *Actin* housekeeping gene for normalization using primers SCL368 and SCL369.

### Disruption of *fksA*, *fksB*, and *fksC* genes

We achieved the partial gene deletions by utilizing our previously developed recyclable marker system [33, 34]. This system utilizes the pyrimidine biosynthesis gene *pyrG* as a selection marker for genetic manipulation due to *Mucor* being intrinsically resistant to most drugs, thus making drug selection impossible. To disrupt our genes of interest we constructed a disruption cassette containing the *pyrG* gene flanked by ∼1k of 5’ and 3’ regions of our gene of interest via overlap PCR. We then inserted the cassette into a TOPO vector. The vector was then propagated in ONESHOT *E. coli* competent cells. We then grew the *E. coli* on selective media containing kanamycin and X-gal for blue white selection. To insert our knockout cassette we used a linearized plasmid what was digested using the SmaI enzyme For *fksA* the 5’ region was amplified primers SL593 and SL543 the 3’ region was amplified with primers SL544 and SL594. For *fksB* the 5’ region was amplified primers SL552 and SL553 the 3’ region was amplified with primers SL554 and SL555. For *fksC* the 5’ region was amplified primers SL314 and SL315 the 3’ region was amplified with primers SL316 and SL317. To further confirm the insertion of our knockout cassette disrupting the *fksA* locus, we produced a 1,880 bp fragment spanning from the 5’ junction to the *pyrG* recyclable marker using primers SL593 and SCL556 and a 2,168 bp fragment spanning the 3’ junction *pyrG* recyclable marker using primers SL594 and SCL557 (Fig 3B). Furthermore, we amplified the open reading frame (ORF) of our mutant strain using primers SL548 and SL549 and produced a 1,090 bp fragment in both the WT and mutant strains (Fig 3B). To confirm the insertion of our knockout cassette disrupting the *fksB* locus, we produced a 1,610 bp fragment spanning from the 5’ junction to the *pyrG* recyclable marker using primers SL552 and SCL556 and a 2,055 bp fragment spanning the 3’ junction *pyrG* recyclable marker using primers SL555 and SCL557 (Fig 3c). Furthermore, we amplified the open reading frame (ORF) of our mutant strain using primers SL558 and SL559 and produced a 896 bp fragment in both the WT and mutant strains (Fig 3d). *Mucor* spores are known to be multinucleated and produce aseptate hyphae, thus the amplification of both the junctions and ORF fragments show that these mutants are heterokaryotic. This results in a partial deletion of these alleles in the mutant strain genome.

### MIC and disc diffusion assays

To measure the extent to which each of the different mutants we generated impacts the susceptibility of *Mucor* to micafungin or caspofungin, we employed a minimum inhibitory concentration (MIC) assay. We inoculated spores of wild-type, *fksA*^wt/Δ^, *fksB*^wt/Δ^, *fksC*Δ, *bycA*Δ, or *bycA*Δ::*cnbR*Δ in RPMI media containing micafungin or caspofungin serially diluted in twelve equal steps ranging from of 256 µg/ml to 0 µg/ml. Minimum inhibitory concentrations were determined according to the Clinical and Laboratory Standards Institute (CLSI) guideline [32]

We utilized a disk diffusion assay to observe a zone of inhibition around paper disks. We plate a lawn of either wild-type, *fksA*^wt/Δ^, *fksB*^wt/Δ^, *fksC*Δ, *bycA*Δ, or *bycA*Δ::*cnbR*Δ on MMC media (pH 4.5) or MMC media (pH 4.5) with 1% activated charcoal to aid as a visual contrast between the media and mycelium. We then place two autoclaved disks; one loaded with micafungin (100 µg) and one with no drug as a control.

### Virulence test

To determine the virulence of the strains we generated, we utilized a wax moth larvae model. For the production of *Mucor* spores, our strains of interest were grown on YPG agar at 26°C for 4 days under light. The spores were washed two times with PBS before injecting a 2μl mixture of 4×10^4^ spores and PBS into the last left proleg of the wax moth larvae host. Differences between the survival curves were evaluated for significance using a Kaplan-Meier test. The experiment was performed on two different occasions with *n* = 15 larvae per group.

We also utilized an immunocompromised murine host intratracheal model. We inoculated mice with the WT strain (R7B) the *bycA*Δ single mutant, the bycA cnbR double mutant, or PBS (mock) via the intratracheal route. Six-week old CF1 Mice were immunocompromised with cortisone acetate (500mg/kg) via the subcutaneous route and cyclophosphamide (250mg/kg) via the intraperitoneal route. Immunosuppression was administered every 5 days starting 2 days before infection while the experiment lasted 12 days (cite). On the day of inoculation, the mice were anesthetized using isoflurane and a mixture of 1×10^6^ spores containing PBS were introduced via the intratracheal route. The treated groups were treated with micafungin (2mg/kg) via the intraperitoneal route on days 0, 5, and 10 We followed with daily monitoring of weight and activity. Differences between the survival curves were evaluated for significance using a Kaplan-Meier test The experiment was performed on two different occasions with *n* = 5 animals per group.

## Statistics

Prism 7 (GraphPad Software Inc.) was used to perform statistical analysis. A *P* value of ≤0.05 was considered significant.

## ACKNOWLEDGEMENTS

We are indebted to xx, yy, zz. This work is supported by a Korean Food Research Institution (KFRI) grant and CTSA/IIMS UT Health pilot grant to S.C.L. S. C. L. holds a Voelcker Fund Young Investigator Award from the Max and Minnie Tomerlin Voelcker Fund. A. G. is supported by the UTSA RISE-PhD program (NIGMS RISE GM60655).

---

**Supplemental Figure 1** Expression of the *fks* genes during various life stages of *Mucor.* We measured the expression of the *fks* genes in the different developmental stages including germination of spores, hyphal formation, and sporulation phages. There were no significant differences in the expression of any *fks* gene regardless of developmental stage.

**Supplemental figure 2.** Confirmation of deletion of *cnaC* gene in *Mucor*. The *cnaC* deletion cassette was generated by an overlap PCR with the recyclable pyrG marker flanked by ∼1 kb fragments of 5’ and 3’ ends of the gene (light green showdown regions). A homologous recombination results in the replacement of cnaC gene with the pyrG marker and the deletion was confirmed by junction PCRs using primers indicated in the figure. The absence of the cnaC ORF was also confirmed by using an ORF specific PCR reaction.

**Supplemental figure 3.** Deletion of the catalytic subunits of calcineurin does not result in susceptibility to echinocandins. A MIC assay was conducted on mutants lacking their respective calcineurin catalytic subunits CnaA, CnaB, or CnaC. The MIC of micafungin for all mutants was >256 µg/ml. The MIC of caspofungin was 256 µg/ml in the *cnaA*Δ mutant and 128 for the *cnaB*Δ and *cnaC*Δ mutants.

**Supplemental figure 4.** PCR confirmation of the disruption of the *fksC* gene. (A) illustration of the *fksC* (upper) and *fksC*::*pyrG* (lower) alleles with 1 kb of the 5’ and 3’ sequences (genes not to scale). (B) Amplification of the 5’ junction (1,849 bp) of the *fksC*::*pyrG* allele and open reading frame (ORF) (1,105 bp) of the *fksC* gene. Lane 1 is the amplification of the 5’ junction in the WT, which should result in no band due to the recyclable marker not being present. Lanes 2 and 3 are the amplification of the 5’ junction in two homokaryotic mutants of *fksC*Δ. Lane 4 is the amplification of the ORF of *fksC* in the WT strain. Lanes 5 and 6 are the amplification of the ORF of *fksC* in two *fksC*Δ mutants. There is no band due to the disruption of the *fksC* gene. (C) Amplification of the 3’ junction in the *fksC*::*pyrG* allele. Lanes 1 and 2 are the amplification of the 5’ junction in two homokaryotic mutants of *fksC*Δ. Lane 3 is the amplification of the 5’ junction in the WT strain, which should result in no band due to the recyclable marker not being present.

**Supplemental figure 5** PCR confirmation of the partial disruption of the *fksA* gene. (A) illustration of the *fksA* gene(upper) and *fksA*::*pyrG* allele (lower) with 1 kb of the 5’ and 3’ sequences (genes not to scale). (B) Amplification of the 5’ junction (1,980 bp) of the *fksA*::*pyrG* allele. Lanes 1 and 2 are the amplification of the 5’ junction of two *fksA*Δ mutants. Lane 3 is the amplification of the 5’Junction of the *fksA*::*pyrG* allele in the WT strain. No amplification is observed in lane 3 due to the recyclable marker not being present. (C) Amplification of the 3’ junction (1,311 bp) of the *fksA*::*pyrG* allele. Lanes 1 and 2 are the amplification of the 3’ junction of two *fksA*Δ mutants. Lane 3 is the amplification of the 3’Junction of the *fksA*::*pyrG* allele in the WT strain. No amplification is observed in lane 3 due to the recyclable marker not being present. (D) Amplification of the open reading frame (ORF) (1,105 bp) of the *fksA* gene. Lanes 1 and 2 are the amplification of the ORF of the *fksA* gene in two *fksA*Δ mutants. Amplification is observed in lanes 1 and 2 due to the genes being partially disrupted resulting in heterokaryotic mutants. Lane 3 is the amplification of the ORF of the *fksA* gene in the WT strain.

**Supplemental figure 6** PCR confirmation of the partial disruption of the *fksB* gene. (A) illustration of the *fksB* gene(upper) and *fksB*::*pyrG* allele (lower) with 1 kb of the 5’ and 3’ sequences (genes not to scale). (B) Amplification of the 5’ junction (1,610 bp) of the *fksB*::*pyrG* allele. Lanes 1 and 2 are the amplification of the 5’ junction of two *fksB*Δ mutants. Lane 3 is the amplification of the 5’Junction of the *fksB*::*pyrG* allele in the WT strain. No amplification is observed in lane 3 due to the recyclable marker not being present. (C) Amplification of the 3’ junction (1,069 bp) of the *fksB*::*pyrG* allele. Lanes 1 and 2 are the amplification of the 3’ junction of two *fksB*Δ mutants. Lane 3 is the amplification of the 3’Junction of the *fksB*::*pyrG* allele in the WT strain. No amplification is observed in lane 3 due to the recyclable marker not being present. (D) Amplification of the open reading frame (ORF) (869 bp) of the *fksB* gene. Lane 1 is the amplification of the ORF of the *fksA* gene in the WT strain. Lanes 2 and 3 are the amplification of the ORF of the *fksB* gene in two *fksB*Δ mutants. Amplification is observed in lanes 2 and 3 due to the genes being partially disrupted resulting in heterokaryotic mutants.

