## supplemental figures for "Duplication and overexpression of the genes encoding a beta 1,3-glucan synthase confer the intrinsic resistance to echinocandin in *Mucor circinelloides*"

Supp figure 1

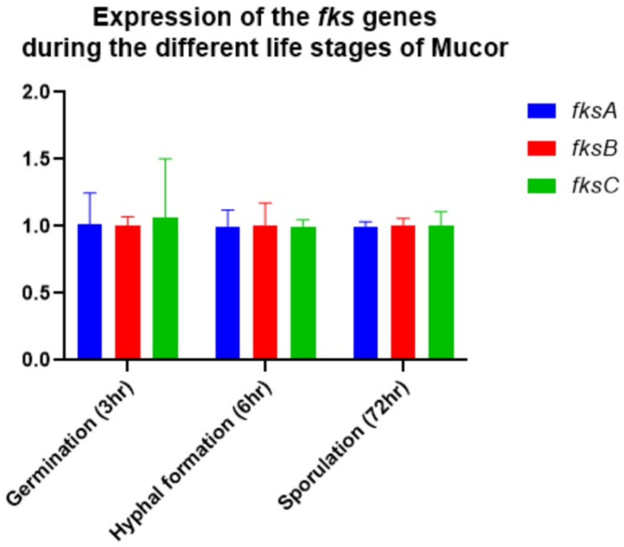

Supp figure 2

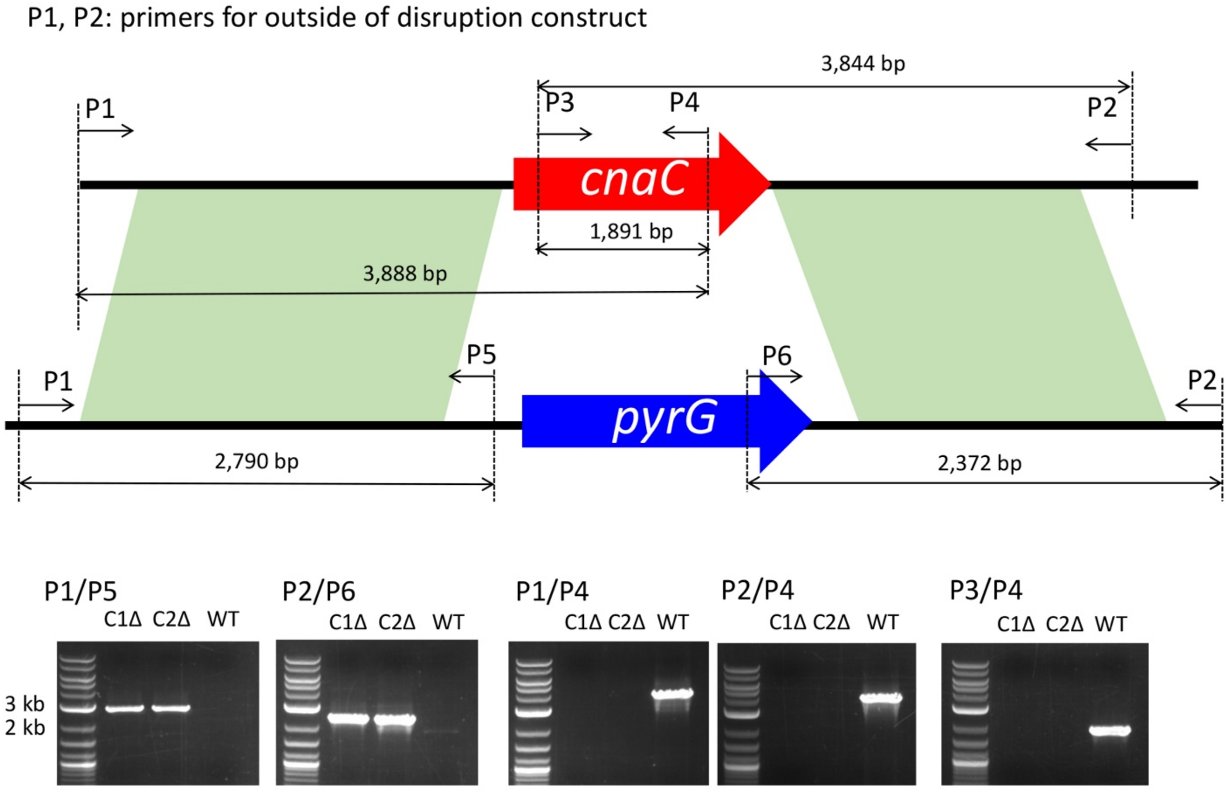


C1Δ: *cnaC1*Δ, C2Δ: *cnaC2*Δ, WT: wild-type

Supp figure 3

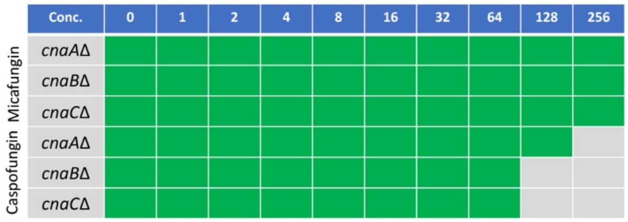

Supp figure 4

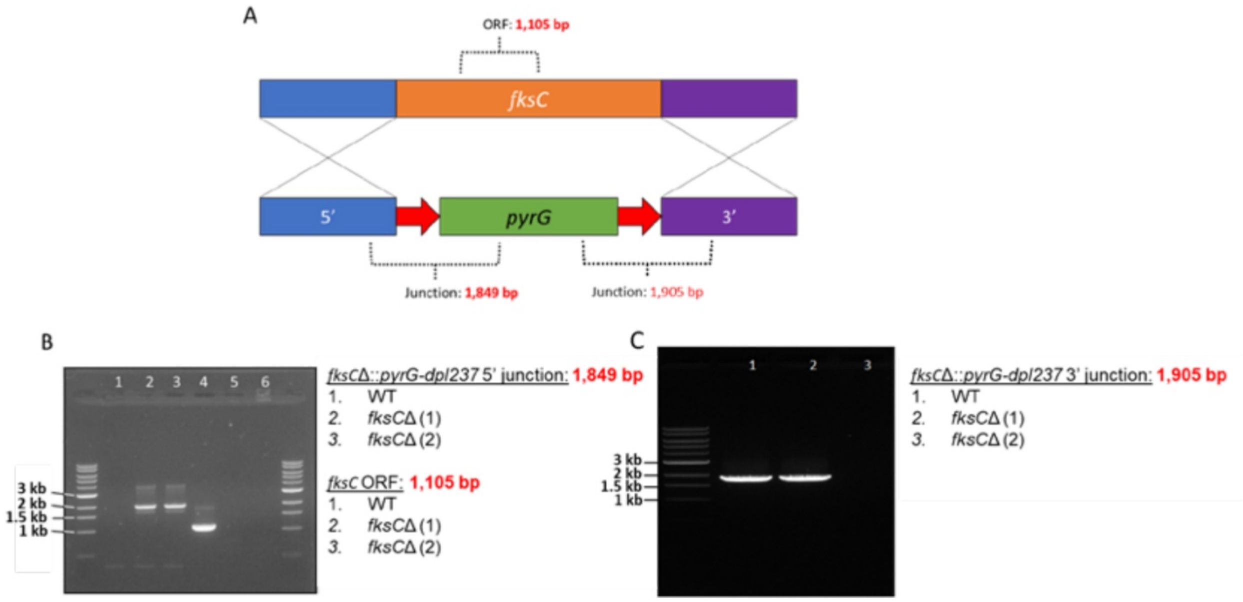

Supp figure 5

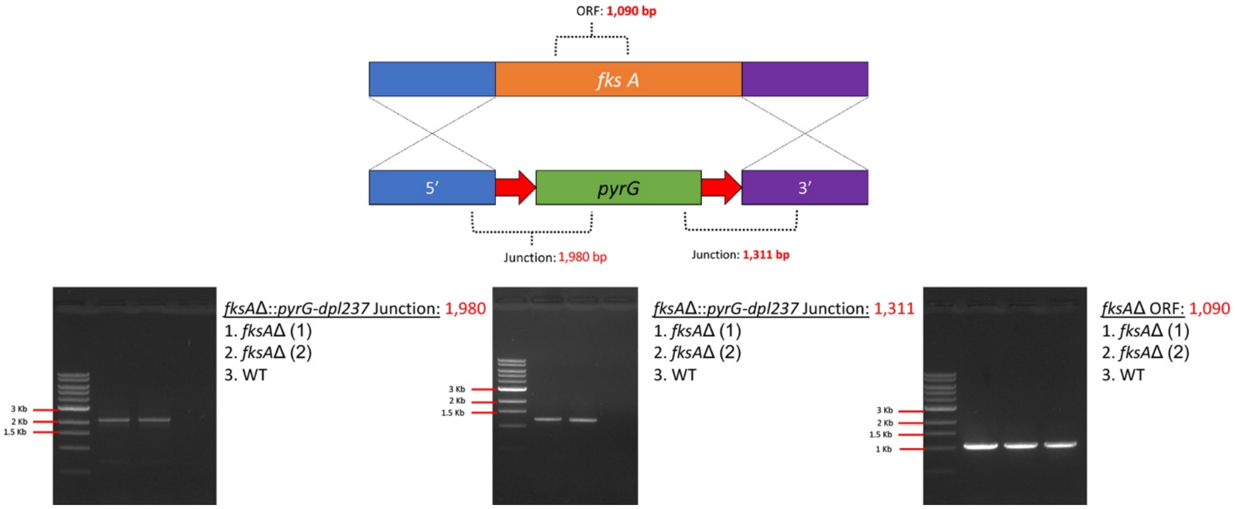

Supp figure 6

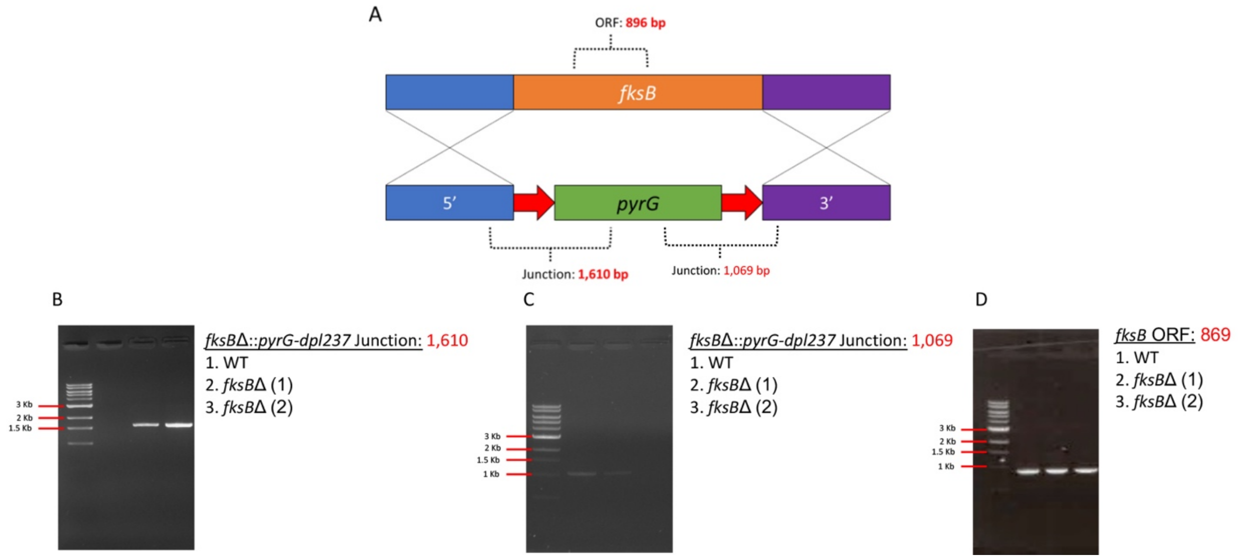
